# Molecular epidemiology of Infectious Spleen and Kidney Necrosis Virus (ISKNV) in Ghanaian cultured tilapia

**DOI:** 10.1101/2022.11.15.516701

**Authors:** Angela Naa Amerley Ayiku, Abigail Akosua Adelani, Patrick Appenteng, Mary Nkansah, Joyce M. Ngoi, Collins Misita Morang’a, Richard Paley, Kofitsyo S. Cudjoe, David Verner-Jeffreys, Peter Kojo Quashie, Samuel Duodu

## Abstract

Infectious Spleen and Kidney Necrosis Virus (ISKNV) is increasingly gaining more attention globally, due to its highly significant economic impact on the aquaculture industry. In late 2018, unusually high levels of mortality (60-90%) was reported in some intensive tilapia cage culture systems in Ghana. Preliminary investigations confirmed the involvement of ISKNV, a viral pathogen noted for fatal systemic infections in many fish species. As a follow-up on the outbreak situation, and post-mass vaccination of affected fish farms, the need to investigate further the molecular epidemiology and phylogeography of the virus across Lake Volta became paramount. In this study, a multiplexed PCR assay and MinION™ nanopore sequencing of the Major Capsid Protein (MCP) were performed to investigate the presence and genotype of ISKNV in tilapia collected from 30 randomly selected farms spread across Lake Volta. Fish with and without clinical signs were included in the molecular detection of the virus from brain, kidney and spleen tissues. ISKNV was detected at 80% prevalence with fry and juvenile fish being most affected. Phylogenetic analysis of the MCP revealed that all 35 isolates from 14 different farms were ISKNV genotype I with near- 100% homology to the 2018 outbreak strain. Vaccination and heat shock treatment; the main specific interventions currently employed to control the viral pathogen have not achieved much success and ISKNV remains a threat to the growth of the aquaculture industry in Ghana. The outcome of this study can be useful in improving fish health management and biosecurity policies in the aquaculture industry.

## 2 INTRODUCTION

Infectious Spleen and Kidney Necrosis virus (ISKNV) is a megalocytivirus known to induce a fatal systemic disease in some aquatic animals resulting in mass mortalities and causing significant economic losses in the aquaculture industry (1–5). Symptomatology of the disease in fish usually includes lethargy, coelomic distension due to ascites, exophthalmia, frayed fin and hemorrhages on body parts (1,6–8). The virus is naked, icosahedral and about 150 nm in diameter (7). The virion core contains a single linear dsDNA molecule of approximately 111kb (9,10).

In recent times the major capsid protein (MCP) gene has been used to classify and assess genetic relationship of unknown megalocytivirus isolates because of its highly conservative DNA sequence (11–14). There are three main genotypic clusters of megalocytiviruses: red sea bream iridovirus (RSIV), first isolated in cultured red sea bream (*Pagrus major*); infectious spleen and kidney necrosis virus (ISKNV), isolated initially from farmed mandarin fish (*Siniperca chuatsi*); and turbot reddish body iridovirus (TRBIV), initially from farmed turbot (*Scophthalmus maxius*) (15–17).

Conventional diagnosis of ISKNV infection is based on histopathological analysis, electron microscopic observation, or immunoassays (1,6,18,19). However, several PCR-based methods such as nested PCR, loop-mediated isothermal amplification (LAMP) and qPCR assays have been established for detecting and quantifying ISKNV infection (2,12,19–22). The host range of ISKNV is broad, predominantly affecting freshwater and brackish water fish species such as tilapia (*Oreochromis* spp.) (1,22,23). The virus was originally identified in a high-mortality outbreak among cultured mandarin fish (*Siniperca chuatsi*) in China and was subsequently detected in other geographies (5,24,25). The export of asymptomatic infected fish has been linked to the spreading of ISKNV widely to other parts of Asia, Europe and Australia; especially in aquaculture facilities (2,9,20,24,26).

Aquaculture of Nile tilapia (*Oreochromis niloticus*) is an emerging industry in Ghana where fish constitutes 50–80% of consumed animal protein. The yearly per capita consumption of tilapia is estimated at 28 kg, significantly higher than most African countries (27–29). Because, Nile tilapia is the dominant and preferable species for fish farming and consumption, the industry employs thousands of people and is critical to livelihoods and the general economy (4,30). In Ghana, from only 2,000 tonnes in 2006, Nile tilapia production soared to over 55,000 tonnes per annum by 2016, with revenues of approximately US$ 200,000,000 (31,32). More than 90% of this production was derived from high stocking-density floating cage systems on Lake Volta, one of the largest artificial lakes in the world (33,34). In 2018 and 2019 production dropped to about 30,000 tonnes per annum representing 46% economic loss due to disease outbreak (35). Infection with ISKNV was detected in few farms in 2018 and 2019 and subsequently, this virus was confirmed to be the major cause of the devastating economic loss to the cage farmers and hatchery operators (36). This was the first reported cases of ISKNV in Africa. To curtail the spread of the virus, the Ghanaian Ministry of Fisheries and Aquaculture carried out a mass immunization of fish farms across the lake using the Aquavac Irido Vaccine, a commercial iridovirus vaccine from Singapore (37). The industry is recovering, but yet to peak to pre-outbreak levels (38).

Since the first report of ISKNV in farmed tilapia in Ghana by Ramírez-Paredes et al (2019), no other studies on ISKNV in the country have been published on the burden, presentation and scale of the disease. There has also not been any report on the outcome of the iridovirus vaccine rollout in Ghana. This comprehensive study developed a molecular assay to detect ISKNV infection accurately from a broad swathe of fish farms on the lake. We performed sequencing and phylogenetic analysis to establish strain relatedness and collected important epidemiological data on disease mortalities and controls measures at the farm level using a detailed questionnaire. We described the wide spread of the virus on the lake and the need to have a better practical control strategies moving forward.

## 3 MATERIALS AND METHODS

### 3.1 Bio-specimen collection

Field sampling was carried out between 25 August and 6 November 2021 in 30 tilapia farms along Lake Volta within Eastern (upstream) and Volta (downstream) regions. This is the hub for intensive floating cage culture systems in the country. To ensure good sampling coverage, fish were collected from only farms in active operation, who gave informed consent (Supplemental data 1), and were adequately spaced out across the study area. The farms sampled included small (n=9), medium (n=12) and large (9) scales of operations. On average, 10 biological samples of fry, fingerlings, grow-out adult fish, brood-stocks and their eggs were collected at each farm depending on the type of facility (i.e., grow-out and/or hatchery) in operation. For farms with both hatchery and grow-out, it was ensured that good representative samples (10-12 fish and egg material) were collected across the entire production cycle. Symptoms indicative of ISKNV infection were recorded for each fish on a separate form (Supplemental data 2). All fish sampled were euthanized with 0.20 mL of clove oil per 500 mL of water for 10 minutes (39). Fish tissues (kidney, brain and spleen, eggs/ova) were preserved in RNAlater, on the field, for subsequent PCR detection of ISKNV in the lab. Ethical approval was acquired from the University of Ghana Institutional Animal Care and Use Committee (UG-IACUC 007/20-21).

### 3.2 Farm interviews and gathering of epidemiological information

A structured electronic questionnaire was developed to collect epidemiological information from all farms visited (Supplemental data 1). This obtained data on farm operations, production data, disease episodes, mortalities, biosecurity, and control measures against ISKNV, as well and other infectious agents. All samples and corresponding questionnaire were linked with a unique farm identification number.

### 3.3 DNA extraction and PCR amplification

Total DNA was extracted from the RNAlater-preserved tissue using the QIAamp DNA Mini Kit (Qiagen); according to the manufacturer’s specification. Briefly, tissue (<25 mg) was removed and washed with sterile 1X PBS (500µl) twice. Glass beads and 100µl of buffer ATL were added to the tissue. The tissue was then homogenized by mechanical disruption and incubated with 20µl of proteinase K at 56°C overnight for 12 hours. After which total nucleic acids were extracted and stored until further use.

Using the extracted DNA as template, detection of ISKNV-like virus from the tissue samples was carried out by multiplex PCR amplification targeting four putative ISKNV genes designed from the nucleotide sequence of ISKNV (GenBank accession number AF371960) using the NCBI Primer Blast tool (Table 1) (10,40). The 25µl PCR mixture consisted of 12.5µl of 2X Go Taq (Hot Start) DNA Polymerase (Invitrogen), 0.5µl of 0.2µM forward and reverse primers for each gene (Table 1), 6.5µl of molecular grade water, and 2µl of DNA template. The PCR reactions included an initial denaturation of 5 min at 95°C; followed by 40 cycles of denaturation at 95°C for 1 min, annealing at 56°C for 1 min, extension at 72°C for 1 min; and a final elongation step at 72°C for 10min. PCR reactions were subsequently visualized by agarose gel electrophoreses. The assay had previously been optimized to detect ISKNV in 200fg/µl of fish-derived nucleic acid (Figure 1). Theoretical specificity of primers used was examined against other genomes by performing basic local alignment searches (41). Aquatic bacterial pathogens (*Streptococcus agalactiae, Escherichia coli, Edwardsiela tarda, Chrysobacterium gambrini, Aeromonas Jandaei Aeromonas veronii, Plesiomonas shigelloides and non-typhoidal Salmonella*) and extracts from tissues of ISKNV uninfected fish were used as reference samples to confirm the analytical specificity of the multiplex PCR assay (data not shown). To test the sensitivity, DNA from three positive ISKNV samples were diluted serially from a starting concentration of 2ng in 8 steps 10-fold dilution. Each dilution step, was used as template in the multiplex PCR as described above. The amplified DNA was analyzed by gel electrophoresis using 1.5% agarose.

**Table 1:**
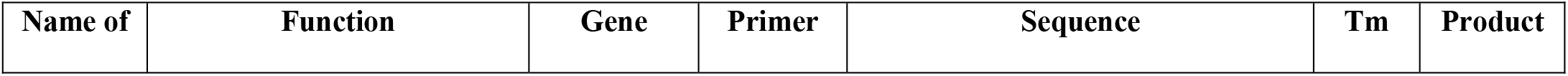
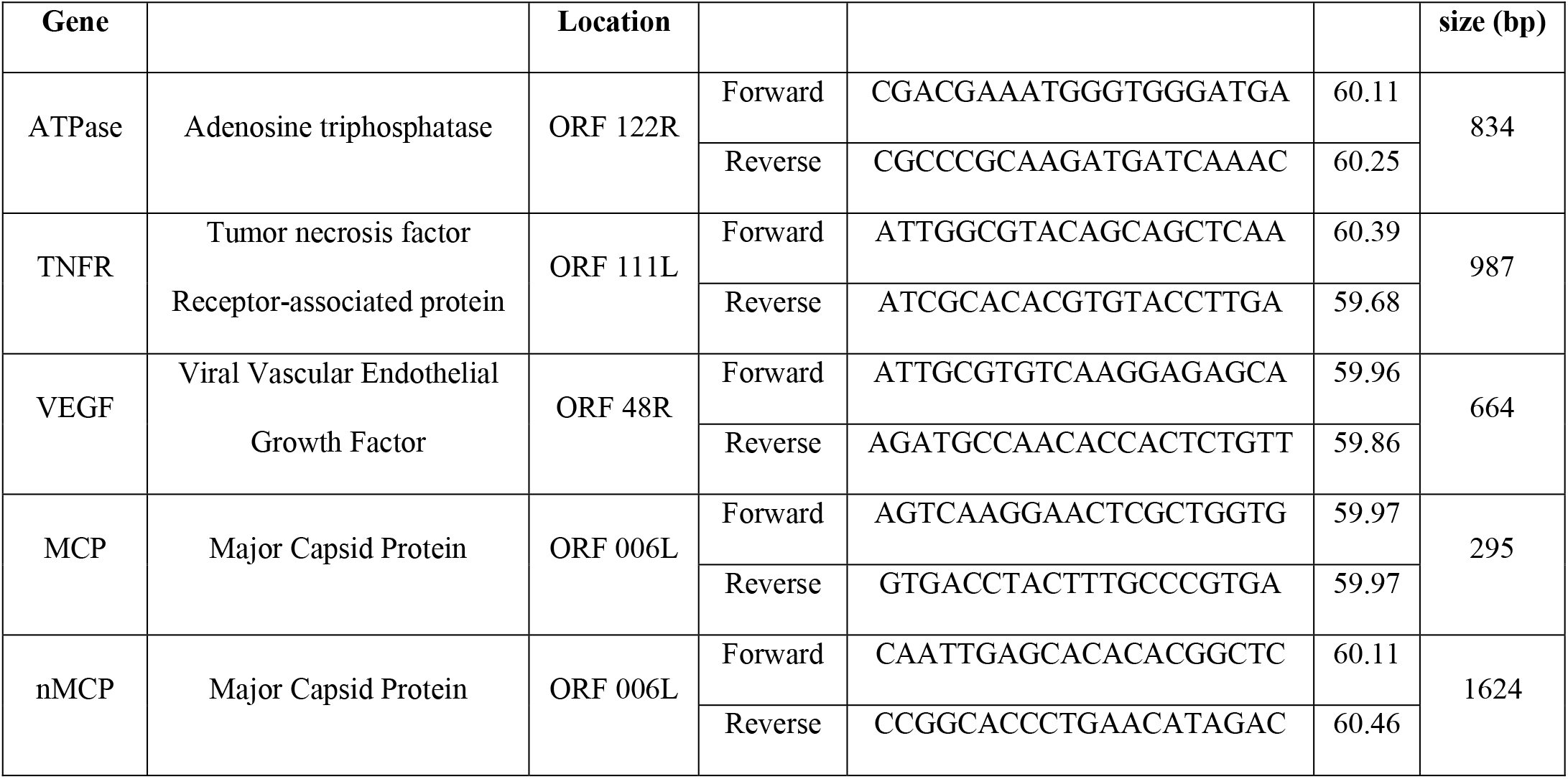
List of primers used for PCR assay

**Figure 1.**
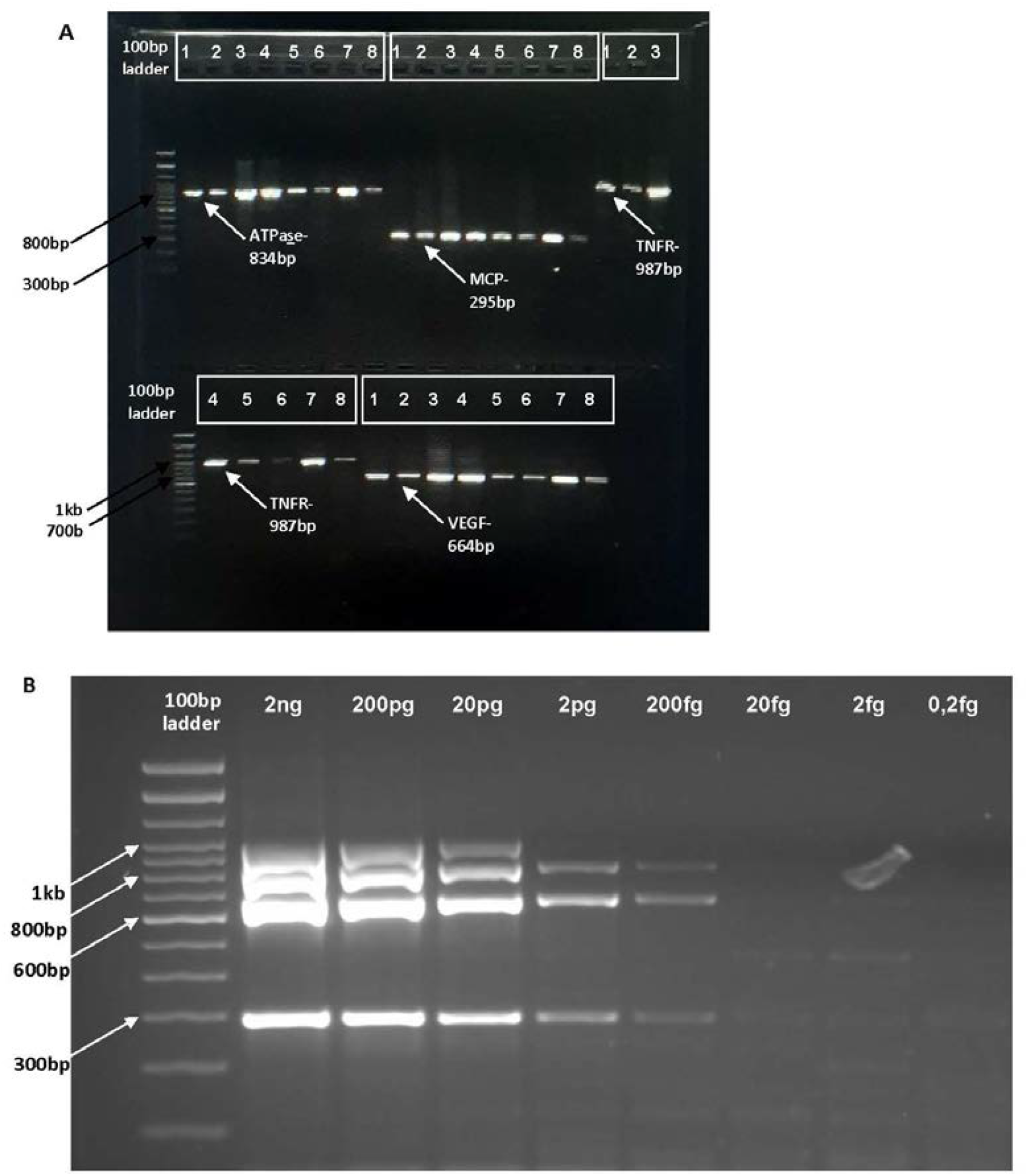
(A) Gel image of showing individual optimization of primers (TNFR (987bp), ATPase (834bp), VEGF (664) and MCP (295)) used for the multiple assay (B) Optimization of multiplex assay.

### 3.4 Sequencing and phylogenetic analysis

The complete MCP gene fragment (1358 bp) was amplified for 40 ISKNV isolates using a different set of primers (nMCP) (Table 1) designed from nucleotide sequence of ISKNV GenBank accession number AF371960. Amplicon sequencing was performed on an Oxford Nanopore™ MinION using a modified protocol based on SARS-CoV-2 whole genome sequencing(42). Sequencing libraries were prepared using the NEBNext Ultra II End Repair/dA-tailing module (New England Biolabs, UK). Amplicons were barcoded using the EXP-NBD196 kit (Oxford Nanopore Technologies, UK). The barcoded amplicons were then pooled, purified with Ampure XP beads (Beckman Coulter) and quantified using the QubitTM DNA HS Assay Kit (Thermo Fisher ScientificTM, USA). About 75 ng of barcoded libraries were ligated to the AMII sequencing adaptors (Oxford Nanopore Technologies, UK) using the Quick ligation kit (New England Biolabs, UK), then finally purified and quantified with Ampure XP beads (Beckman Coulter) and QubitTM DNA HS Assay Kit (Thermo Fisher ScientificTM, USA), respectively. Approximately 20 ng of the purified adaptor-ligated libraries were subsequently loaded on an R9.4.1 flow cell (FLO-MIN106) to be sequenced using a MinION Mk1b device (Oxford Nanopore Technologies, UK). Base-calling and demultiplexing of MinION Fast5 files were achieved using Guppy (Version: v5.0.7) (43,44). Alignments and statistics to the reference sequence (AF371960) was done using epi2me-labs/wf-alignment (Version. v0.1.8), followed by a consensus sequence generation using NGSpeciesID (Version - v0.1.2.1) (45–47). Finally sequence alignment and visualization of alignment were performed using MAFFT (Version: v7.394) and MEGA 11, respectively (48–50).

Nucleotide sequence analysis of ISKNV MCP gene fragments from various isolates and geographical locations were compared with 13 known Megalocytivirus sequences retrieved from GenBank. Multiple sequence alignment was performed with CLUSTALW (51). A phylogenetic tree based on the MCP nucleotide sequences was constructed using the Neighbor-Joining method (52). The evolutionary distances were computed using the Maximum Composite Likelihood method and were in the units of the number of base substitutions per site. Evolutionary analyses were conducted in MEGA11 (48). Sequences were submitted to GenBank with Accession numbers OP689616-50.

## 4 RESULTS

Across the thirty sampled farms, common external abnormalities or disease signs observed in sampled fish included lethargy, cloudy/opaque eyes, exophthalmia, loss of eyes/ skin discoloration, loss of scales, skin erosions, distended abdomen, fin rot and hemorrhages on body parts (Figure 2). Fish with no observable symptoms such as the above and appeared healthy yet tested positive for ISKNV were subsequently referred to as asymptomatic.

**Figure 2.**
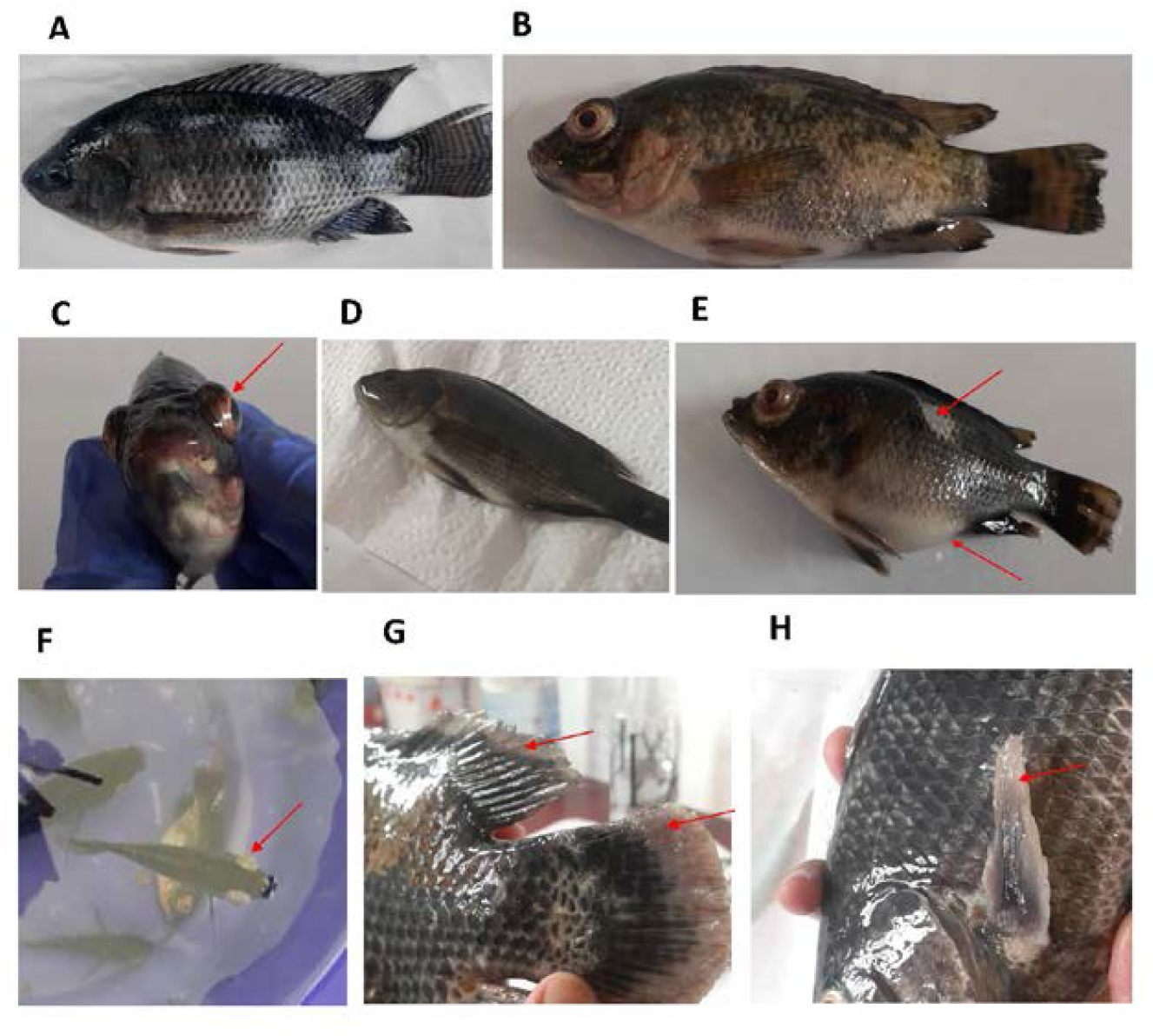
Some abnormalities (disease symptoms) observed in the sampled fish; (A) asymptomatic, (B) discoloration, hemorrhaging and frayed fin, (C) Exophthalmia, (D) excess mucus on the skin, (E) Ascites & loss of scales, (F). opaque eyes, (G) eroded caudal fin, (H) eroded pectoral fin.

### 4.1 Molecular detection of ISKNV

Samples were considered positive if they showed bands for all four putative genomic regions (Figure 3). Of the 30 farms screened, ISKNV was detected in 24 farms, representing 80% (Figures 4 & 5) and 110 out of 254 fish tissue samples assayed from these farms were positive (43.3%). A majority of the farms screened were on the Eastern and Western banks. Here, ISKNV was detected in 21 of the 24 farms (87.5%). In the Volta region 3 out of 7 farms (42.8%) showed positivity. At the time of sampling most of these farms were closing down or demonstrated significantly reduced scales of production Generally, ISKNV was predominantly detected in fry and juvenile fish (≤20g) (65.4%) as compared to adult/grow-out fish (>20g) (Table 2). In addition, 6.25% (2 out of 32) of eggs screened were positive and only one broodstock out the 24 sampled was positive. Of all the fish found infected, 47.4% were asymptomatic and appeared healthy.

**Figure 3.**
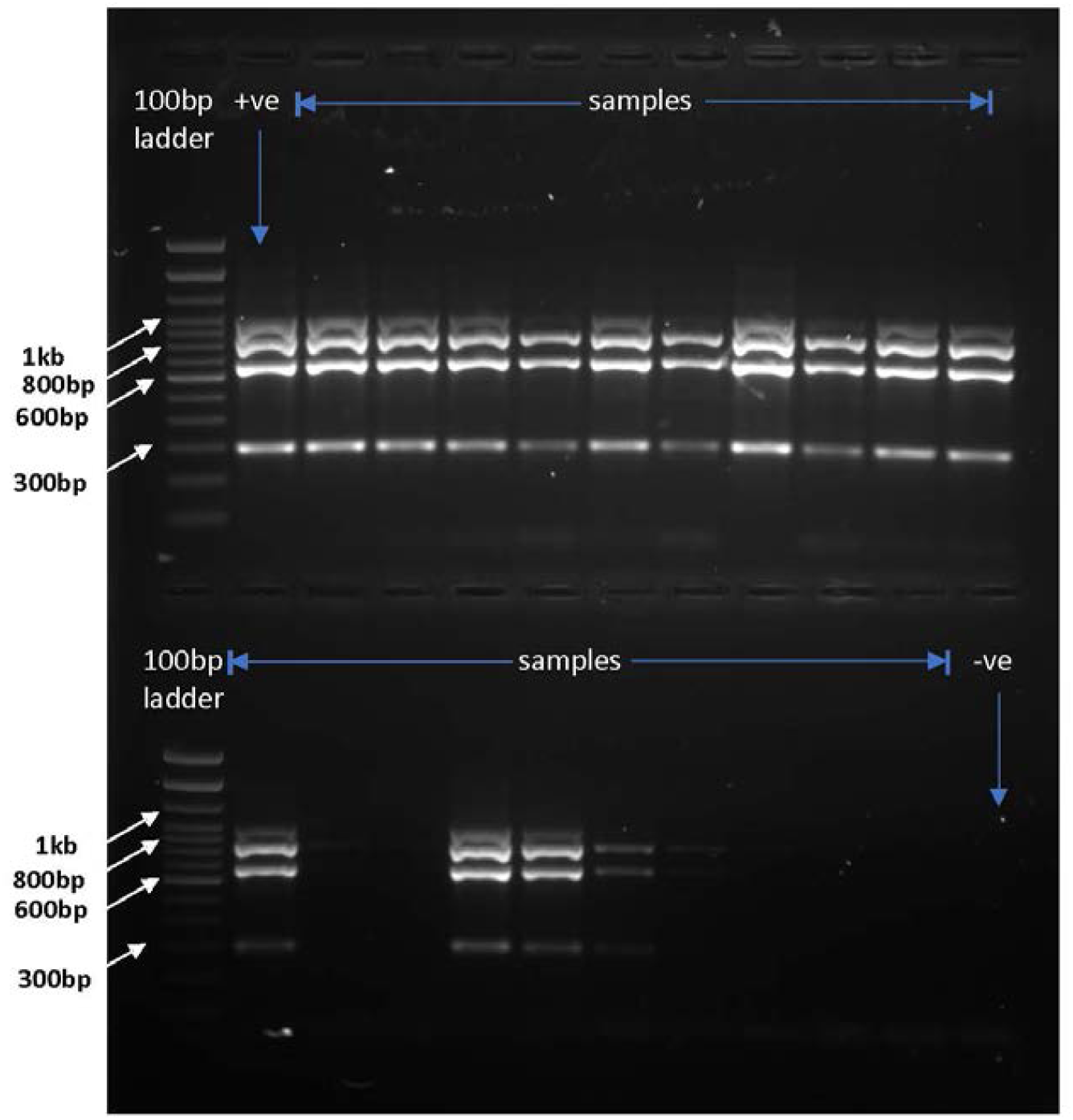
Representative gel image of samples screened using multiplex PCR.

**Figure 4.**
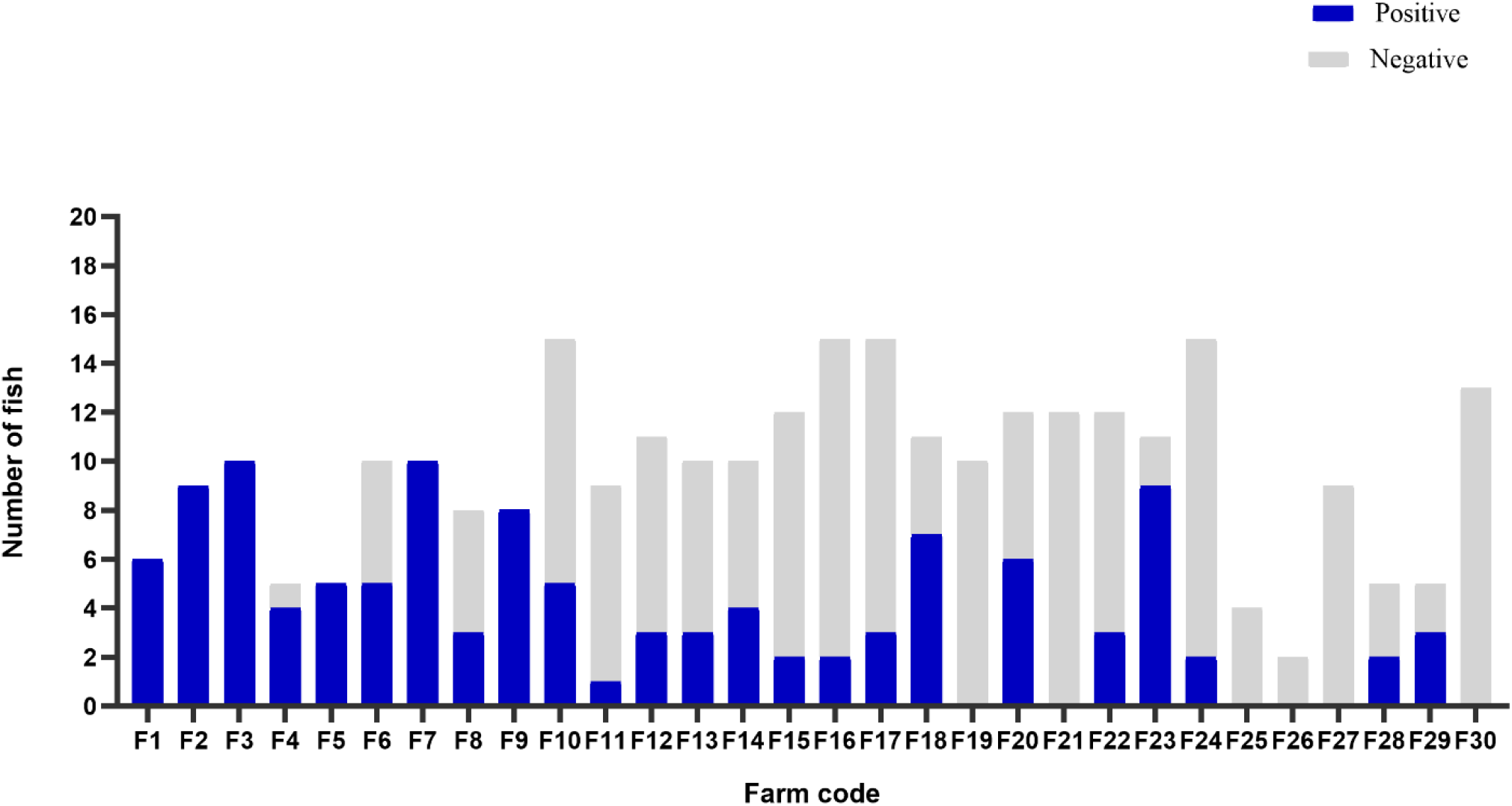
Graph showing PCR test results of sampled fish per farm

**Figure 5.**
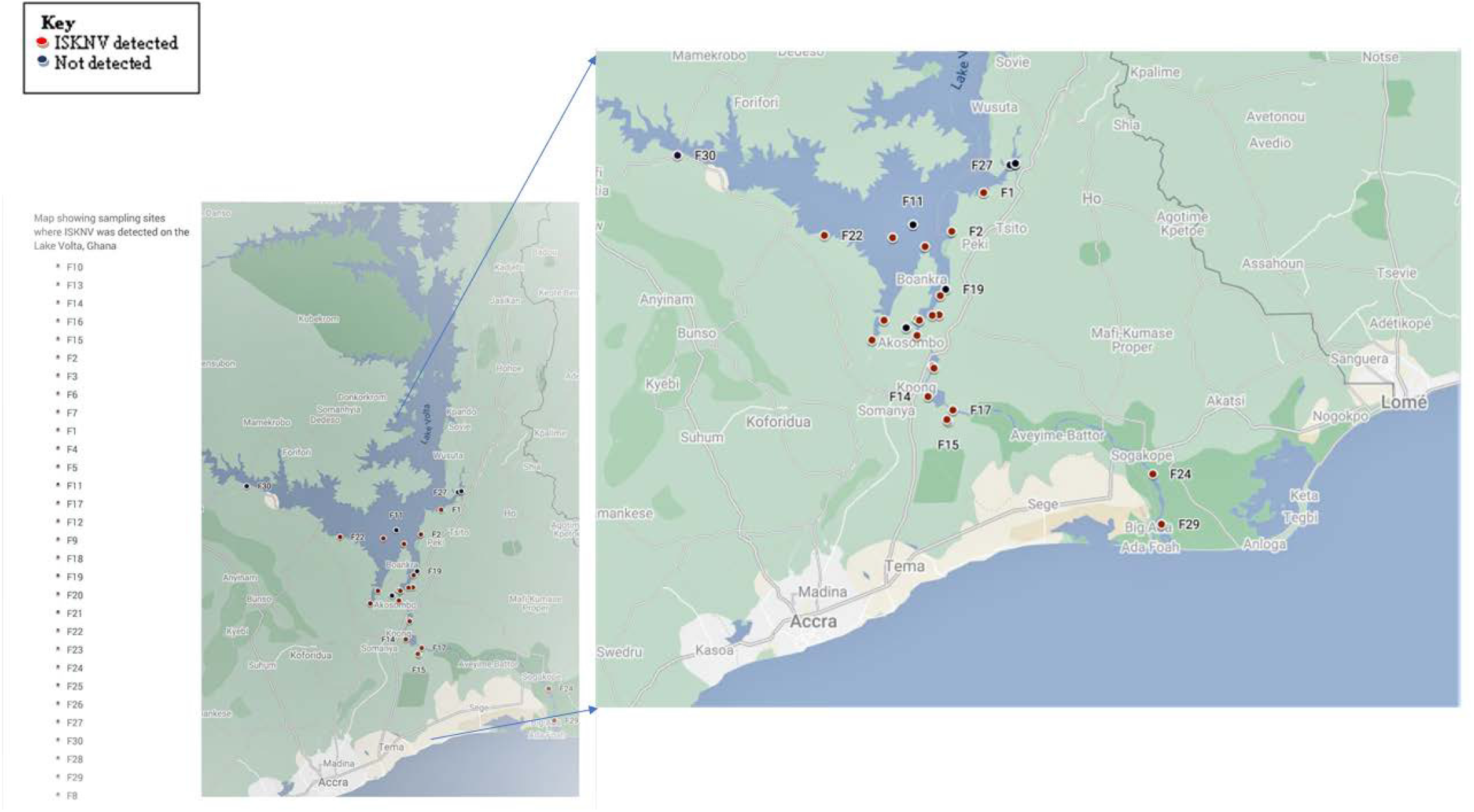
Map showing the distribution of ISKNV infected farms on Lake Volta

**Table 2:**
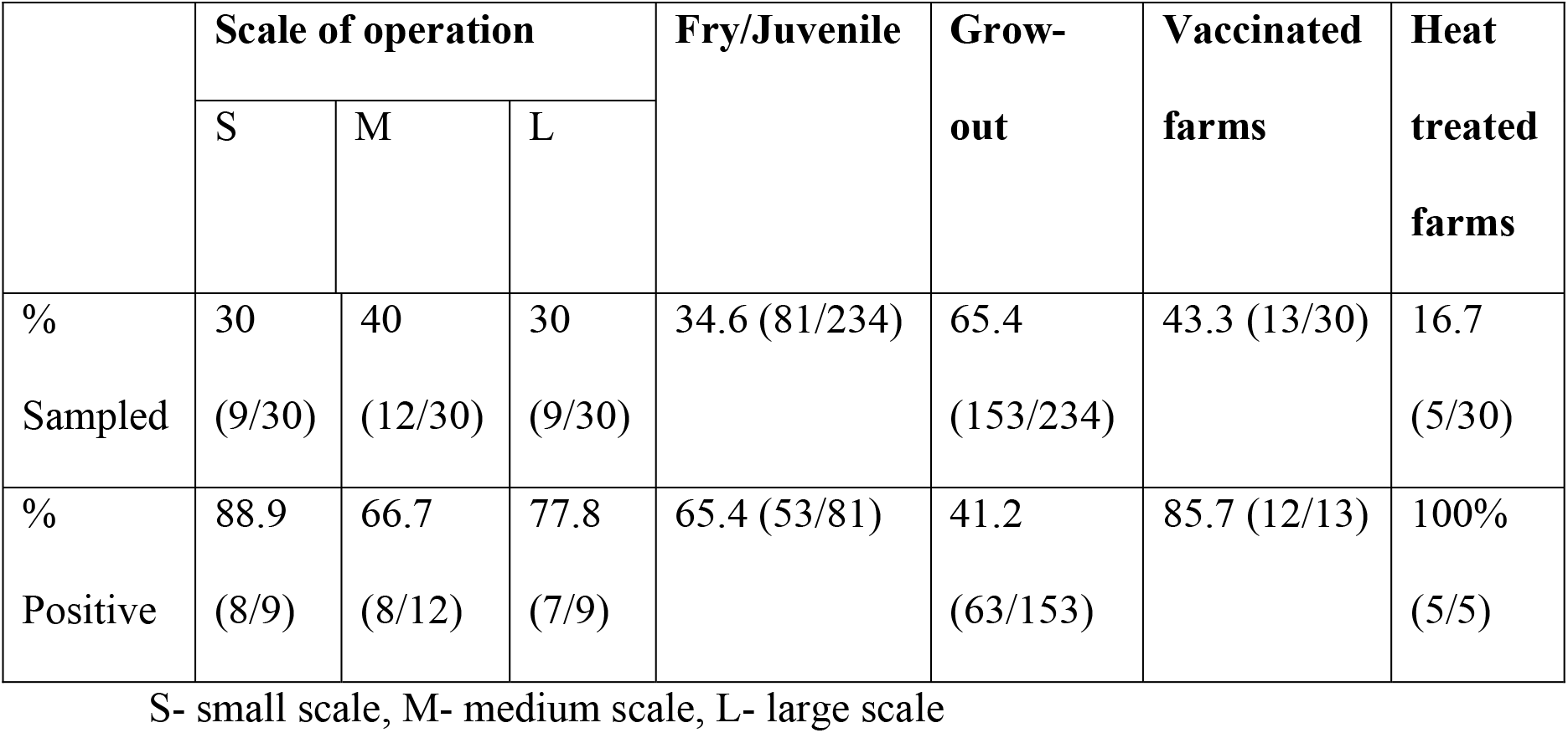
Representation of proportion of samples collected for each category listed and the percentage found to have been infected.

### 4.2 Responses of farm interviews

Most of the studied farms (60%, n=18) had been in operation between 1 to 10 years and the remaining 40% (n=12) had been in operation for more than 10 years. Reported daily pre-outbreak mortality ranged from 0.1 to 65%, with a median of 1% and a mean of 6%. Only four farms reported a baseline mortality above 10%. Twenty-eight farms (93.33%) reported having experienced significant disease outbreak in the past five years with high mortality (22 farms, range 40-95%) and large financial losses (median 70%, range 75 – 90%). There were also reports of significantly higher mortality in fingerlings than juveniles and grow-outs. Currently, the reported average mortality is 40%.

To study transmission patterns, the source of broodstock and fingerlings was recorded (Figure 6). Most farms were outsourcing either both broodstock and fingerlings or only broodstock (fingerlings produced in house) and a few outsourced only fingerlings. Farmer-reported observations included one or more of the following clinical signs: unusual swimming, skin nodules, loss of eyes, darkening of eyes, bulging of eyes, loss of scales, excess mucous on skin, distended abdomen, discoloration/darkened skin, weight loss, skin lesions, whitening of mouth part, frayed fins, eroded fin, fin rot and fish lice. Interestingly, 10 farms reported observable changes in water color prior to disease events with changes in measured parameters. Vaccination and heat-shock treatment are intended to prime the immune system of the fish to produce helpful immune component that increases the chances of surviving a natural encounter with the virus (23,53). These were the most reported ISKNV-specific interventions adopted by the farmers. Salt treatment, reduced feeding, use of immuno-stimulants, herbal treatment, reduced stocking density, in-house fingerling production, net cleaning, removal of diseased fish and antibiotic treatment were less frequently mentioned. In addition, one farm derives its fingerlings in-house by crossbreeding the Akosombo strain, the national accredited farmed strain with wild tilapia species from Lake Volta, as an intervention to the viral disease. Thirteen farms out of those sampled had received the Aquavac Iridovirus vaccine once. The vaccine was administered intra-peritoneally to fish weighing 3 grams and above after anesthetization with clove oil. Farms that received the commercial vaccine rolled out by the government, reported an improvement in survival rate but still had mortalities rates above 40%. For heat-shock treatment, ISKNV exposed fish were gradually introduced to temperatures 10°C above the optimal growing temperature (27-30°C) for 30 mins (54). This was repeated over a period of 6 days. When asked how interventions could be improved, key inputs included; improved diagnostics, regulation of fingerling production and sales, fingerling health certification, regulation of chemicals, reduced stocking density, improved vaccines, vaccination strategies and heat treatment regimen.

**Figure 6.**
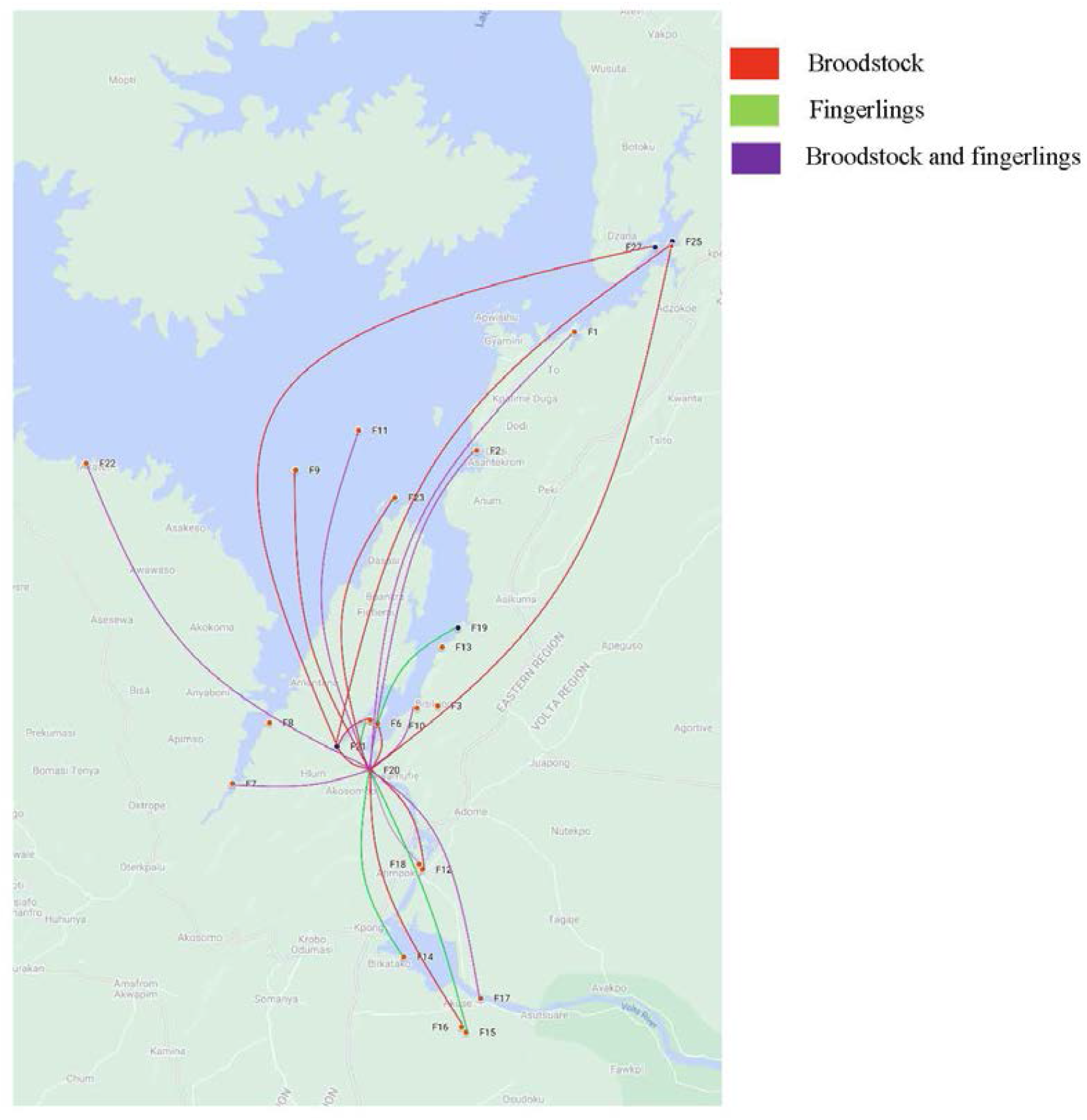
Map showing the movement of fingerlings and brood stock across Lake Volta from farm to farm

### 4.3 Sequencing and phylogenetic analysis of MCP gene

The PCR assay using the nMCP primer set to amplify the full MCP genomic region yielded expected PCR products (1634bp) (Supplemental data 3) from 35 of the multiplex assays confirmed isolates. The optimal tree generated from phylogenetic analysis is shown in figure 7. The bootstrap values at 100 replicates are shown next to the major nodes [2]. This analysis involved 48 nucleotide sequences. All positions containing gaps and missing data were eliminated (complete deletion option). There were 1360 positions in the final dataset. Sequence analysis of these new amplicons showed that samples were identical to each other with the exception of F22S1. The viruses identified from 2018 were reported to show 100% homology with the ISKNV Clade 1 reference sequence AF371960 (36). Based on the phylogenetics, all successfully sequenced isolates also clustered with AF371960 including the marginally diverged virus (Figure 7).

**Figure 7.**
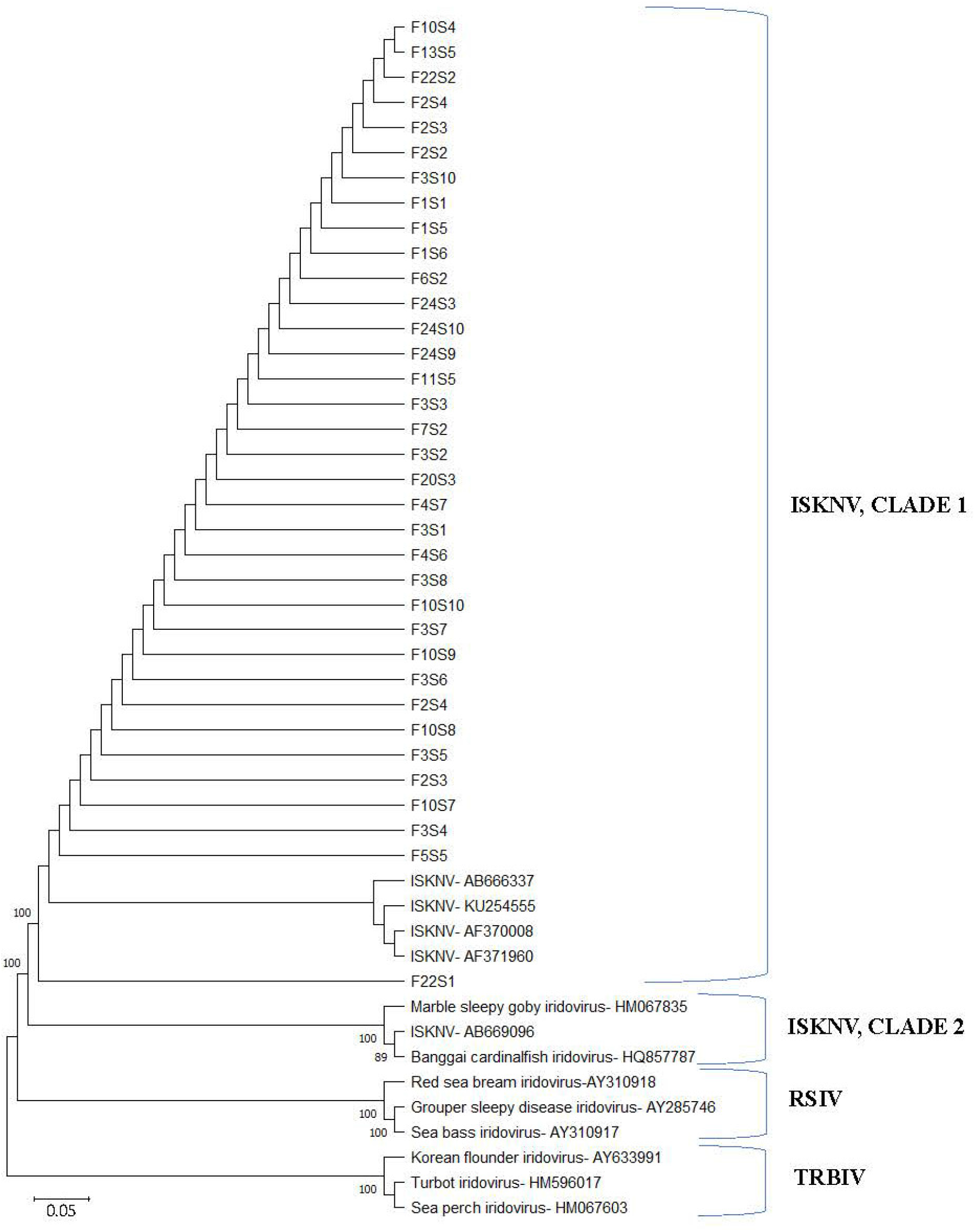
Phylogenetic tree of 35 sequenced samples and 13 reference sequences from the GenBank using the Neighbor-Joining method conducted in MEGA11. The bootstrap values for 100 replicates are shown at major nodes. The evolutionary distances were computed using the Maximum Composite Likelihood method and the scale bar is for the number of base substitutions per site.

## 5 DISCUSSION

Megalocytiviruses are well known for their broad host range including both ornamental and food fish, and their ability of more than one genotype infecting the same host species (12,55). The MCP gene is most suitable for analysis of phylogeny among iridoviruses, due to its relatively conservative nucleotide sequences (12,56). Megalocytivirus show high nucleotide and amino acid sequence identities and ISKNV have been found to have comparatively lower genetic variation and much more unique biological features (24). Molecular screening conducted in this study to detect specifically the virus in tilapia, amplified all four putative genes (MCP, VEGF, TNFR and ATPase) in a multiplex PCR assay. The full MCP genomic region was amplified to further confirm the viral pathogen and for the test of phylogeny. The sequence analysis revealed that all isolates based on the MCP were ISKNV genotype clade 1. Expectedly, the sequence identity of the isolates infers that these isolates were of the same strain and that no variation of sequences had occurred over the three years’ period since the virus was first detected in the country (Figure 8). The sequences clustered with ISKNV sequence (AF371960.1) which exhibited high homology with the 2018 ISKNV isolate from Ghana. This indicates that the circulating iridovirus causing high mortalities across the farms on Lake Volta is ISKNV clade 1 and is very likely still descended from the original strain if new introductions are responsible for current mortalities. Given the scale of coverage of this study, we can tentatively say that Ghana is no longer experiencing an ISKNV outbreak but rather an epidemic in farmed tilapia. This information can inform vaccine design and deployment strategies.

**Figure 8.**
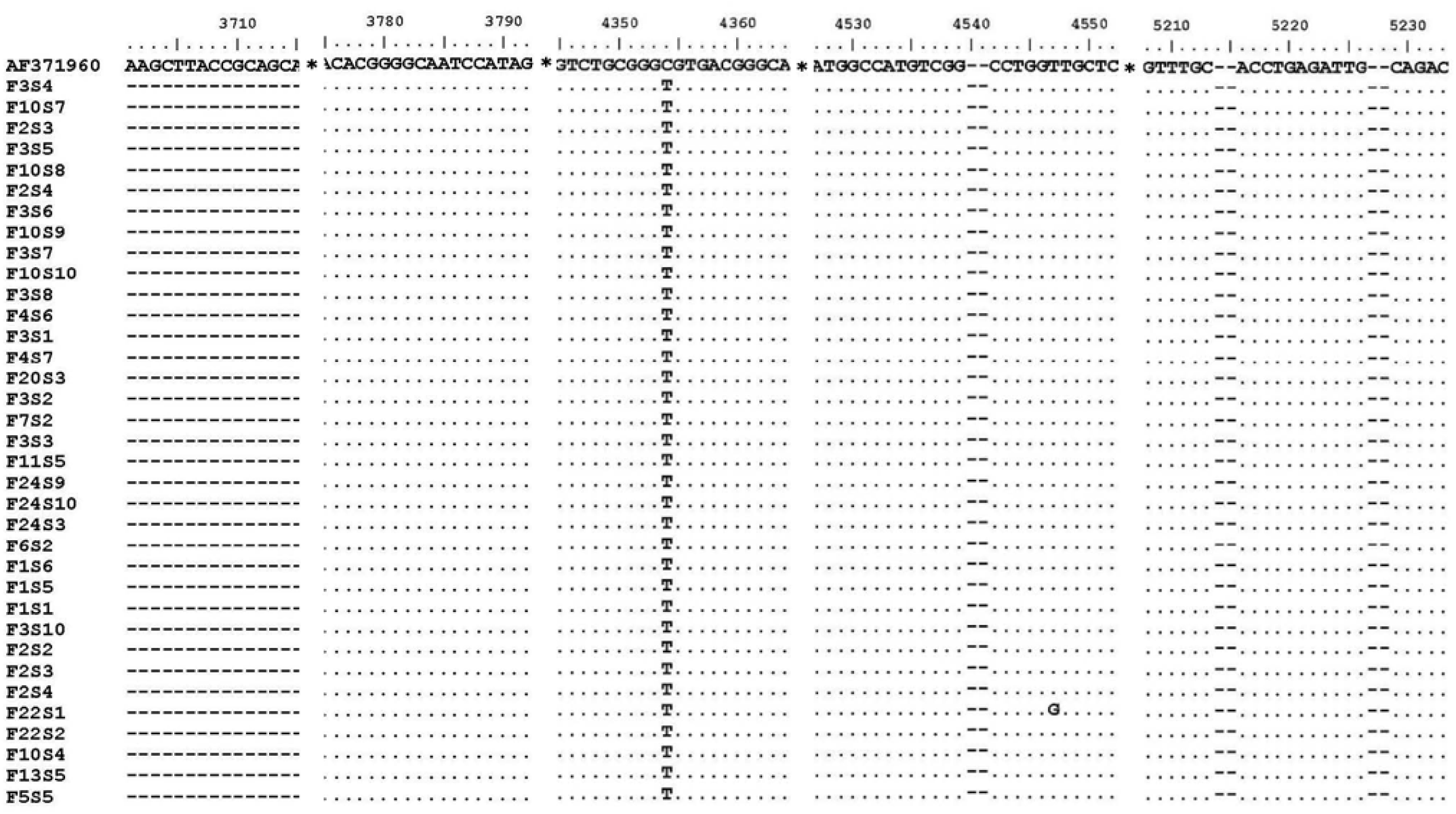
Multiple alignment of sequences from study samples with the reference sequence (AF371960).

ISKNV was highly prevalent amongst all farm types and across operational scales, with an apparent overall farm-level prevalence of 80% (24 out of 30 farms). The high prevalence and wide geographical distribution of the virus within the study area suggests significant horizontal transmission between farms. Common horizontal transmission routes in aquaculture includes live infected fish movements, fomite contact between farms and local spread through water contact (7). One of the farms with significant high positivity in their samples (54.5%) also supplies broodstock and/or fingerlings to 17 other farms in this survey. Two farms which received only broodstock were negative and the other 15 farms were all positive. Several farms reported having biosecurity measures in place. However, effective biosecurity measures were found to be largely absent, with only 3 farms having biosecurity physical barriers (vehicle and footwear dips, signpost) mounted at the farm gate and other strategic entry points during farm visits. This may increase the likelihood of inadvertent horizontal transmission between farms. Also, almost one half (47.4%) of the sampled fish that tested positive for ISKNV, showed no external symptoms (Figure 2), making the transfer of supposedly healthy fish to other farms more likely. ISKNV was detected in eggs (6.25%) and broodstock (4.17%) in the current study, suggesting possible rare occurrence of vertical transmission (3).While there have not been any publications describing true vertical transmission of ISKNV, its detection in broodstock and eggs highlight the importance of enforcing biosecurity measures and testing at hatcheries to prevent spread from such facilities. With a limited number of operational hatcheries in Ghana, many farmers still depend on a few hatcheries to source their fingerlings thereby promoting large-scale live fish movements over large geographical areas.

Most ISKNV-negative farms were distant from neighboring farms or located further offshore, which may indicate that these farms are exposed to a lower disease pressure or may easier implement effective biosecurity measures preventing the introduction of the virus. Farm-level routine disease prevention practices including disinfection (use of salt or other disinfectant), access restriction, staff biosecurity training, heat treatment, reduced feeding, herbal remedies, removal of dead and diseased fish, net cleaning, not sharing equipment and some biosecurity measures were reported. During the mass vaccination in 2019, 13 farms received the commercial Iridovirus vaccine (37). All these farms were included in this study. Twelve of these farms were found to be still positive for ISKNV with high mortality possibly indicating an ineffective response to the vaccine. Indeed, most farmers could only partially testify to the vaccine efficacy as mortality rates had reduced slightly but still remained comparatively very high at the time of sampling. Moreover, the possibility of vertical transmission threatens the vaccination programme as fingerlings may already be infected before vaccination. It would be prudent to also focus vaccination on the few hatcheries available in the country, to ensure that juvenile fish have a better chance at survival once introduced into the Lake. Also, vaccine efficacy studies must be carried out to ensure that the vaccines, are yielding the expected response. Although some farmers considered heat-shock treatment an effective pathogen control strategy, a well-structured system for hyperthermia treatment and an understanding of the mechanism of action is required to evaluate its beneficial role in disease management properly.

Concerning management interventions, the wide and prolonged use of unregulated broad-spectrum antibiotic could introduce resistant bacterial strains in the aquatic environment. This unfortunately poses a threat to terrestrial animals as well as humans due to introduction of multi-resistant pathogenic bacteria and antibiotic toxicity from consumption of these fish (57,58). Maintaining good water quality was critical for increased survival of tilapia against ISKNV infection, especially in the hatchery. With an abundant resource such as the Lake Volta, the aquaculture industry has great potential of meeting the high protein demand in the country and beyond (27,59).

The findings from this study suggest that ISKNV introduction to Ghanaian tilapia aquaculture was likely through a single source and has reached epidemic status, possibly through the transfer of juvenile fish between farms on the Volta Lake. Novel measures such as heat shock, especially when combined with vaccination appeared to have some success against mortality, indicating the possibility of managing the ISKNV disease situation with the appropriate control strategies.

Introduction of disease diagnostic laboratories that carry out regular monitoring of production facilities, keeping of proper production data and reporting of unusual mortality coupled with strict enforcement and adherence to biosecurity measures can provide early warnings and protect the industry from surprise mass mortalities. Furthermore, the high prevalence of ISKNV in the vaccinated farms necessitate the need for due diligence in the procurement and administration of vaccination programs. Commercial vaccines must demonstrate efficacy and meet some basic regulatory requirements.

## 6 CREDIT AUTHORSHIP CONTRIBUTION STATEMENT

Angela Naa Amerley Ayiku: Methodology, Investigation, Data curation and analysis, Writing – original draft. Abigail Akosua Adelani: Methodology, Investigation, Data curation. Patrick Appenteng: Methodology. Mary Nkansah: Methodology. Joyce M. Ngoi: Methodology. Collins Misita Morang’a: Methodology, Data curation. Richard Paley: Conceptualization, Supervision, Funding acquisition. Kofitsyo S. Cudjoe: Methodology, Funding acquisition. David Verner- Jeffreys: Conceptualization, Supervision, Funding acquisition. Peter Kojo Quashie: Methodology, Supervision, Funding acquisition, Writing- review & editing. Samuel Duodu: Conceptualization, Methodology, Investigation, project administration, Funding acquisition, Supervision, Writing – review & editing.

## 8 ACKNOWLEDGMENTS

All farmers who participated in this study are thanked for their cooperation and provision of information. The Fisheries Commission of Ghana is appreciated for their support. We thank Hannah Segbefia, Deborah Mettle and Kukua Thompson for their helpful assistance.

